# Peripheral B cell populations tune spontaneous neuronal activity in the uninjured hippocampus after stroke

**DOI:** 10.64898/2026.03.02.709034

**Authors:** Thomas A. Ujas, Navid S. Tavakoli, Pavel Yanev, Vanessa O. Torres, Jadwiga Turchan-Cholewo, Xiangmei Kong, Erik J. Plautz, Adam D. Bachstetter, Nancy L. Monson, Lenora J. Volk, Pavel I. Ortinski, Ann M. Stowe

## Abstract

B cells infiltrate the contralesional hippocampus following stroke, but whether lymphocytes modulate post-stroke plasticity and neuronal network function remains unknown. To identify immune cell mechanism(s) supporting remote plasticity, we examined the impact of B cell depletion on synaptic and neuronal activity in the hippocampal circuit following stroke. Basal synaptic transmission in the contralesional dentate gyrus (DG) following a stroke in adult male mice was decreased with B cell depletion. Expanding our studies to encompass the CA1 and DG regions of the hippocampal circuit in male and female mice of different ages, we utilized synapsin-Cre/GCaMP6s mice to visualize spontaneous calcium activity during a 3-week B cell depletion with and without prior stroke. Systemic B cell depletion in the absence of injury altered neuronal activity in the DG, suggesting a novel neuromodulatory role for circulating immune cells. Stroke increased Ca^2+^ transient amplitudes in the contralesional DG and CA1, with B cell depletion again reducing DG amplitudes while increasing the frequency of Ca^2+^ transients. Robust linear regression revealed significant main effects and higher-order interactions (depletion×sex×age×injury), including increased Ca^2+^ transient amplitudes in older post-stroke mice lowered by systemic B cell depletion, though overall the DG appears more sensitive to modulation versus CA1. These results suggest that circulating B cells can tune hippocampal network activity dependent on age, sex, and the presence of brain injury. The selective vulnerability of the DG to depletion-age-injury interactions opens an avenue for future studies on region-specific neuroimmune crosstalk during post-stroke cognitive recovery.

**Presubmission inquiry for *Neuron*:** We uncover crucial insights on the capacity of circulating B cells to directly modulate hippocampal network activity, showing that B cells are not just passive players, but active neuromodulators whose effects are dependent on sex, age, and stroke injury status. In fact, B cells are central players to functional recovery whose evolving role shifts over time, from acutely beneficial and neurotrophic to chronically maladaptive, depending on timing, context, and responding B cell subset. Our study demonstrates a mechanistic link between systemic immune modulation and neuronal calcium activity. This integrative perspective aligns with Neuron’s mission to publish studies that link cellular processes to systems-level functions. These novel findings also add to a more unified model of neuro-immune interactions that highlights how immunotherapies could be harnessed to improve neuronal function during stroke and aging, with several FDA-approved immunotherapeutics available to modulate systemic adaptive immune responses.

## Introduction

Our prior work found that B cells do not impact hippocampal neurogenesis under physiologic conditions but support stroke-induced ipsilesional neurogenesis in young male mice.^1^ Furthermore, the absence of B cells negatively impacted hippocampal- and amygdalar-mediated cognitive function in uninjured mice, inducing anxiety and memory deficits that were exacerbated after stroke.^1^ These studies suggest B cells support post-stroke neuronal function, but they did not directly test effects on neuronal plasticity. B cells infiltrate remote brain regions, including the contralesional hippocampus, with sustained post-stroke B cell depletion reducing ipsilesional hippocampal neurogenesis and compromising neuronal cell survival.^1^ These effects were concomitant with delayed motor recovery, impaired spatial memory, and increased anxiety through 8 weeks post-stroke compared to wildtype (WT) controls with endogenous B cell levels intact.^1^

Neurogenesis, as identified by doublecortin (DCX)^+^ neuroblasts, is induced after stroke in both the ipsi- and contralesional dentate gyrus (DG),^2^ though with more DCX-expressing cells detected ipsilesionally.^3^ While the ipsilesional hippocampus is damaged by ischemic stroke as it is often within the infarct and peri-infarct regions, the uninjured (i.e. contralesional) side also undergoes selective changes. Contralesional CA1 pyramidal neurons exhibit higher neuronal complexity, reduced spine density, and changes in spine volume and length, suggesting structural plasticity and synaptic remodeling secondary to compensation for lost input.^4^ In another rodent model of transient ischemia, CA1 pyramidal cell death was observed at 4 days post-stroke, with neuronal degeneration observed up to 45 days.^5^ General hippocampal volume loss 6 weeks post-stroke is clinically related to cognitive decline and post-stroke dementia,^6,7^ while a clinical fMRI study found that decreased functional connectivity between the hippocampus and the inferior parietal lobule was associated with impaired memory function.^8^ This study also found that functional connectivity was increased between the hippocampus and the cerebellum, thus driving a potential compensatory mechanism after stroke^8^ and highlighting reorganization within the uninjured hemisphere critical for functional recovery.

To examine how B cells influence contralesional hippocampal function after stroke, we used two complementary approaches: slice electrophysiology and *ex vivo* calcium imaging. Slice electrophysiology provides high-resolution measurements of synaptic strength and transmission properties at individual synapses. Calcium imaging, using GCaMP6s (GCaMP is a genetically encoded calcium indicator composed of green fluorescent protein (GFP), calcium-binding protein calmodulin (CaM) and M13 peptide sequence that interacts with CaM), allows us to visualize activity patterns across large populations of neurons simultaneously, revealing network-level dynamics. Together, these methods reveal both the cellular mechanisms (synaptic changes) and functional consequences (network activity patterns) underlying hippocampal recovery after stroke. Specifically, we utilized a Synapsin-Cre x GCaMP6s transgenic mouse line, enabling neuron-specific expression of the GCaMP6s indicator. GCaMP6s has previously been used to study post-stroke neuronal activity, confirming utility in these studies.^9^ In the present study, our data show the capacity of circulating B cells to modulate hippocampal network activity, suggesting B cells are not just passive players but active neuromodulators whose effects are dependent on sex, age, and stroke injury status.

## Methods

All mice were housed with *ad libitum* access to food and water and were maintained in humidity and temperature-controlled colony room on a 12-hour light/dark cycle (lights on at 7:00am). All study procedures were approved by the Institutional Animal Care and Use Committee of the University of Kentucky and UT Southwestern Medical Center and applied in accordance with the NIH Guide for the Care and Use of Laboratory Animals.

### Electrophysiology in acute hippocampal slices of human (h)CD20 transgenic mice

Male mice (3-8 months old, n=6/group) were assigned to each of the 4 experimental groups (i.e., Rituximab-treated uninjured or post-stroke, hCD20^-^ (WT) or hCD20^+^ (B cell-depleted) mice). 100µg of Rituximab was given via intraperitoneal injection to hCD20-TamCre^+/+^ and hCD20-TamCre^−/−^ (littermate controls) mice for three consecutive days prior to inducing tMCAo.^10^ Following tMCAo, an additional 100µg of Rituximab was administered weekly to target the turnover of B cells. Within the uninjured and post-stroke WT groups were 2 hCD20^+^ mice that were unsuccessfully B cell depleted and thus considered “WT” mice. Mice were anesthetized with isoflurane prior to rapid decapitation. 390µm transverse hippocampal slices (2-4 slices per mouse) were prepared using a Leica VT1200S vibratome following dissection of the hippocampus in ice cold oxygenated (95% O_2_/5% CO2) dissection buffer containing the following (in mM): 2.6 KCl, 1.25 NaH_2_PO_4_, 26 NaHCO_3_, 211 sucrose, 10 glucose, 0.75 CaCl_2_, 7MgCl_2_. Slices were recovered for at least 2 hours in 30°C artificial cerebral spinal fluid (aCSF) containing the following (in mM): 125 NaCl, 3.25 KCl, 25 NaHCO_3_,1.25 NaH_2_PO_4_·H_2_O, 11 glucose, 2 CaCl_2_, 1 MgCl_2_. Field excitatory postsynaptic potentials (fEPSPs) were digitally evoked (Cygnus Instruments, Model PG400A) by stimulating the medial perforant path with a bipolar platinum/iridium concentric electrode (FHC, Bowdoinham, ME) above the upper blade of the DG cell body layer. A glass recording electrode filled with aCSF was positioned in the cell body layer approximately 200µm from the stimulating electrode. The signals were amplified by a differential amplifier (Model 1800; A-M Systems), digitized using Axon Instruments Digidata 1550A (Molecular Devices), and monitored using Clampex software (Molecular Devices). Recording aCSF and temperature were identical to recovery conditions, with a flow rate of ∼3mL/minute. Input-output curves were obtained by evoking 2-4 responses per stimulation intensity, increasing from 10µA in 5μA increments. Analysis was performed in Clampfit, with all traces from one stimulation intensity averaged to produce a single fEPSP slope or population spike amplitude measurement.

### tMCAo

Thirty-one animals underwent a 45-minute transient middle cerebral artery occlusion (tMCAo). Anesthesia was induced (2% isoflurane in O_2_ (0.3 mL/min) and N_2_O (0.7 mL/min)). General analgesic (Buprenorphine Sustained-Release, 0.05mg/kg) was administered subcutaneously and a local anesthetic (Marcaine, 0.08% in sterile saline) administered. The mouse was placed on a heating pad and a rectal probe checked animal temperature. A small incision into the temporalis muscle allowed a 0.5 mm flexible laser doppler probe (TSI, St. Paul, MN, USA) to record baseline cerebral blood flow (CBF). The left common carotid artery (CCA) was isolated and permanently ligated. A vertical incision into the CCA was made for the introduction of a 6-0 monofilament (Doccol, Sharon, MA, USA) and gently moved through the CCA into the internal carotid artery (ICA) and advanced ∼10 mm distal to the carotid bifurcation, past the origin of the MCA. CBF was monitored to show a reduction of at least 70% of the pre-ischemic value. Upon successful occlusion, the filament was secured using a temporary suture threaded under the CCA and remained for 45-min prior to removal from anesthesia. At 40 min., the animal was again anesthetized, and CBF measured to confirm ischemia, the filament removed, and reperfusion allowed. CBF verified that reperfusion took place. The threshold for a successful tMCAo reperfusion was to measure greater than 50% of pre-stroke CBF value. The wound was sutured, and the mice received a subcutaneous dose of saline for hydration (0.5 mL of 0.9% Sodium Chloride). Animals were kept on a heating pad at 25-28°C and given food gel in ventilated cages with *ad libitum* access to food and water in normal housing conditions. Animals were checked twice daily for 7 days post-surgery. Of the 31 animals that underwent tMCAo surgery, 8 mice were excluded, including 6 that died within 7 days post-surgery (79% tMCAo survival success rate; 33% B cell depleted), and 2 animals were excluded due to no GCaMP6s expression at time of experiment. Of the 28 non-tMCAo control animals, 2 died before data were acquired (100% B cell depleted), and 1 animal did not have GCaMP6s expression.

### B cell depletion

For B cell depletion, a mouse anti-human CD20 antibody was used (Genentech, San Francisco, CA, USA) while control animals received a mouse IgG2a isotype control antibody (GP120:9674, Genentech). Anti-CD20 antibody stock concentration was 23.12 mg/mL and diluted down to 1 mg/mL in sterile PBS. The IgG2a control antibody stock concentration was 12 mg/mL and diluted to 1 mg/mL in sterile PBS. 100 µL was delivered via intraperitoneal injection to each mouse, once per day for three days. Booster injections were given every 6-7 days to maintain depletion levels, and depletion was maintained for 3 weeks (19-22 days) prior to calcium imaging. For animals that underwent tMCAo, the 3 serial injections were given prior to surgery to not interfere with the post-op recovery.

### B cell depletion Testing via Flow Cytometry

To confirm the efficacy of our B cell depletion protocol, we used 8 mice to compare the anti-CD20 antibody to an IgG2a control antibody; 4 mice were treated with anti-CD20 and 4 mice were treated with an IgG2a control antibody for a period of 3 weeks. Spleens were extracted, processed and analyzed using flow cytometry to identify immune cell populations in the two groups.

Splenocytes were added to 1 mL of 1X PBS for washing prior to staining for Fluorescence-Activated Cell Sorting (FACS) (438 g, 10 min, acceleration 9, brake 9, 4°C). Cells were incubated in Ghost Dye 780 (APC-Cy7) for live/dead classification for 30 minutes, in the dark at 4°C. After, the cells were washed 2x with FACS buffer. An FcR blocker was used to prevent unspecific staining (Miltenyi Biotec, 130-92-575) for 5 min at room temperature. Without washing, cells were stained with the following: CD45 (BUV-805), CD3 (BV-480), CD4 (BUV-396), CD8β (BV-650), CD19 (PE-Cy7), CD11b (BB-515), Gr-1 (BV-421), NK.1.1 (BV-711). After washing 2x in FACS buffer, the cells were then fixed in 1%PFA/0.1% EDTA for 30 minutes, in the dark at 4°C. Cells were washed, resuspended with 300µL of FACS buffer. Collection of samples occurred on a BD FACS Symphony A3 flow cytometer, with 5 lasers (red, blue, ultraviolet, yellow, green). Compensation of cells occurred using unstained cells and beads (Invitrogen UltraComp eBeads, 01-222-42) for single color controls. Supplemental Figure 1 shows a comparison of the CD19^+^CD3^-^ populations in an IgG2A control treated animal (left) and an anti-CD20 treated animal (right). The mice that received anti-CD20 antibody had a group average of 2.42% B cells compared to 54.5% in our IgG2A control-treated animals, proving that our anti-CD20 method of B cell depletion was successful.

### MRI protocol and procedure

Magnetic resonance imaging (MRI) was performed *in vivo* 6–9 days post-stroke to evaluate lesion extent prior to calcium imaging and electrophysiological recordings. Imaging was conducted using a 7 Tesla BioSpec small animal MRI scanner (Bruker BioSpin, Billerica, MA, USA) equipped with a 16 cm horizontal bore and a 400 mT/m gradient system. Mice were anesthetized (2% isoflurane/95%O_2_/5%CO_2_) and positioned supine on a heated animal cradle. The head was fixed using a bite bar and ear bars to minimize motion artifacts. Physiological parameters, including heart rate and oxygen saturation, were continuously monitored and maintained within normal physiological ranges. Core body temperature was monitored via a rectal probe and maintained at 37.5°C using a feedback-controlled heating system. T₂ relaxation maps were generated from multi-echo images acquired using a TurboRARE sequence with the following parameters: repetition time (TR) = 2750 ms, first echo time (TE₁) = 33 ms, echo spacing (ΔTE) = 8.25 ms, and 8 echoes. Twenty-three contiguous slices were collected with a slice thickness of 0.5 mm, a field-of-view (FOV) of 18 × 15 mm², and an in-plane matrix size of 256 × 256 (frequency × phase encoding), with 12 signal averages. T₂ maps were obtained on a voxel-by-voxel level, using an in-house MATLAB 2023a script (MathWorks, Natick, MA, USA). Infarct volumes were calculated by manual tracing of hyperintense regions on T₂-weighted images using ImageJ 1.53 (ImageJ, National Institutes of Health, Bethesda, MD, USA).^11^ Total lesion volume was estimated using the trapezoidal rule, which integrates the cross-sectional infarct areas across slices.^11–13^ Specifically, the infarct areas delineated on each slice were summed and multiplied by the inter-slice distance (0.5 mm), yielding an approximation of the total infarct volume.

### Widefield calcium imaging in GCAMP6S mice

Genotype-confirmed Synapsin-Cre x GCaMP6s transgenic mice were used to visualize and measure neuronal activity in real time. In total, 67 animals were used for the experiment.

### Calcium imaging solutions

Ice-cold aCSF cutting solution contained (in mM): 93 NMDG, 2.5 KCl, 1.25 NaH_2_PO_4_, 30 NaHCO_3_, 20 HEPES, 25 glucose, 5 sodium ascorbate, 2 thiourea, 3 sodium pyruvate, 10 MgSO_4_·7H_2_O, and 0.5 CaCl_2_·2H_2_O (adjusted to pH = 7.4 with NaOH, 300–310 mOsm). This solution was continuously oxygenated with 95%O_2_/5%CO_2._ Slices were allowed to recover in aCSF cutting solution at 34-36°C for 30 min during which increasing volumes of 2 M NaCl (up to a total of 1 mL NaCl/37.5 mL aCSF) were added as previously described.^14^ After recovery, the slices were then transferred to a recording aCSF solution maintained at room temperature. The recording aCSF solution contained (in mM): 130 NaCl, 3 KCl, 1.25 NaH_2_PO_4_, 26 NaHCO_3_, 10 Glucose, 1 MgCl_2_, and 2 CaCl_2_ (pH = 7.2-7.4, when saturated with 95%O_2_/5%CO_2_).

### Sectioning for *ex vivo* Calcium imaging

The mice were deeply anesthetized with isoflurane and euthanized by decapitation. The brain was quickly extracted and kept in a chilled, continuously oxygenated (95%O_2_/5%CO_2_) artificial cerebrospinal fluid (aCSF) cutting solution. Coronal brain slices (300 μm-thick) were prepared with a vibratome (VT1200S; Leica Microsystems, Wetzlar, Germany). During sectioning, each slice was transferred to an oxygenated aCSF recording solution. Slices were maintained, in the dark, in the recording solution at room temperature (while oxygenated) until transferred to the imaging chamber.

### Generating Calcium imaging data

The recording chamber was continuously perfused (1-2 mL/min) with oxygenated recording aCSF warmed to 32 ± 1°C using an automatic temperature controller (Warner Instruments). Spontaneous fluorescence of neuronal GCaMP6s was captured by concatenating five one-minute videos at 30-second inter-video intervals to minimize photobleaching and binned at 512 × 512 pixels, using the ORCA-Flash 4.0 (V2) digital camera (Hamamatsu, Shizuoka, Japan) under wide-field illumination with an LED light source (X-Cite XLED1, Excelitas Technologies, Miamisburg, OH, USA). The videos were acquired through a 40x objective (0.65 μm/pixel) at 25 frames per second. Imaging was alternated between the dentate gyrus (DG) and CA1 regions of the hippocampus.

### Offline analysis of calcium imaging data

Analysis of videos were performed using Fiji (ImageJ; NIH, Bethesda, MD, USA) and MATLAB (The Mathworks, Inc., Natick, MA, USA). Somatic regions of cells showing spontaneous activity (active cells) were outlined as regions of interest (ROIs). These ROIs, and the fluorescence events within them, were detected using the wavelet ridgewalking method.^15^ Somatic calcium event amplitudes, durations, and frequencies were extracted by custom-written MATLAB scripts and averaged for each identified ROI. A total of 12,065 cell regions of interest were recorded across all animals and median values for the CA1 and DG fluorescence in each animal were evaluated. A small number of amplitude measurements were markedly elevated. Specifically, 28 individual cells exhibited amplitudes >100 among 12,065 recorded cells. The corresponding imaging videos were manually reviewed to confirm that these values reflected true events rather than motion or segmentation artifacts. Additionally, we calculated composite indices of calcium activity for inclusion in the supplemental analyses. The Calcium Activity Index was computed as the product of transient amplitude x duration x event frequency. Calcium Load was computed as amplitude x duration.^16^ These are reported in the Supplementary Materials in **Figure 3A-D**.

### Statistical analysis

Statistical analyses were performed using GraphPad Prism version 10.0 (GraphPad Software, San Diego, CA, USA) as well as JMP Pro version 17.2 (SAS Institute Inc, Cary, NC, USA). Descriptive statistics, including means, medians, and standard deviations were calculated to summarize data characteristics. When running statistics on the number of cells recorded, we ran Welch’s t-test (assuming unequal variances) when comparing two groups. When running statistics on the number of cells recorded with age as a continuous variable, we fitted models using robust regression with the Cauchy M-estimator in JMP Pro. Normality was assessed using the Shapiro-Wilk test. For data comparing two groups, Mann-Whitney U tests were performed, while for non-normally distributed data comparing more than one group, the Kruskal-Wallis test was used, followed by Dunn’s multiple comparisons. In JMP Pro, we used the Box-Cox transformation to normalize data and stabilize variance in the dataset. This was performed on amplitude and duration data so that it could be used in a statistical fit model. Using JMP Pro, we performed multiple linear regression using the *Fit Model* platform. Specifically, the Standard Least Squares personality was used. Model terms were selected based on experimental design and biological relevance. We evaluated the main effects of injury, depletion, sex, and age (and their interactions) across hippocampal regions. Effect leverage plots were then used to evaluate the influence of each predictor variable on the model fit to further identify effects and key observations. Additionally, when analyzing for the relationship between infarct volume and age we performed a bivariate fit and used Robust Cauchy Fit estimation method to account for outliers and non-normality. This was also used when looking at the relationship between amplitude and age. It should be noted that JMP Robust Chi-square statistic (used in the Cauchy Fit) is a distinct test that uses quantitative/continuous variables and tests whether a robust regression model fits better than the mean of the response.

## Results

### B cells support synaptic transmission in the contralesional dentate gyrus

To determine if B cells may function to regulate synaptic transmission, we measured input-output curves in the hippocampal DG from uninjured mice or in the contralesional DG 2 weeks following a transient middle cerebral artery occlusion (tMCAo; **Figure 1A**) in adult male mice. Rituximab successfully depleted B cells in mice expressing human CD20 (hCD20^+^)^1^ when compared to hCD20^-^ (i.e., wild-type; WT) littermate controls (F_(1,20)_=6.48, p=0.02; **Figure 1B**). T2-weighted magnetic resonance imaging (MRI) was performed 6 days after stroke to assess the extent of stroke injury prior to electrophysiological recordings and revealed no significant difference in the infarct volumes of WT and B cell-depleted mice (**Figure 1C,D**). Stroke significantly decreased synaptic transmission in B cell-depleted (F_(1, 21)_=4.96, p=0.037), but not WT mice (**Figure 1E,F**), suggesting that the absence of B cells renders DG synapses vulnerable to dysregulation following a stroke. Interestingly, we observed a trend toward increased synaptic transmission following B cell depletion in uninjured mice (p=0.05; **Figure. 1G**), as evidence that B cells may modulate synaptic transmission or neuronal excitability in the healthy hippocampus. Neither B cell depletion nor stroke affected paired-pulse ratios (**Supplemental Fig. 2**), indicating that synaptic release probability is unaffected by the presence of B cells. These data suggest the capacity for direct modulation of synaptic transmission by peripheral B cells independent of their canonical immune functions.

**Figure 1.**
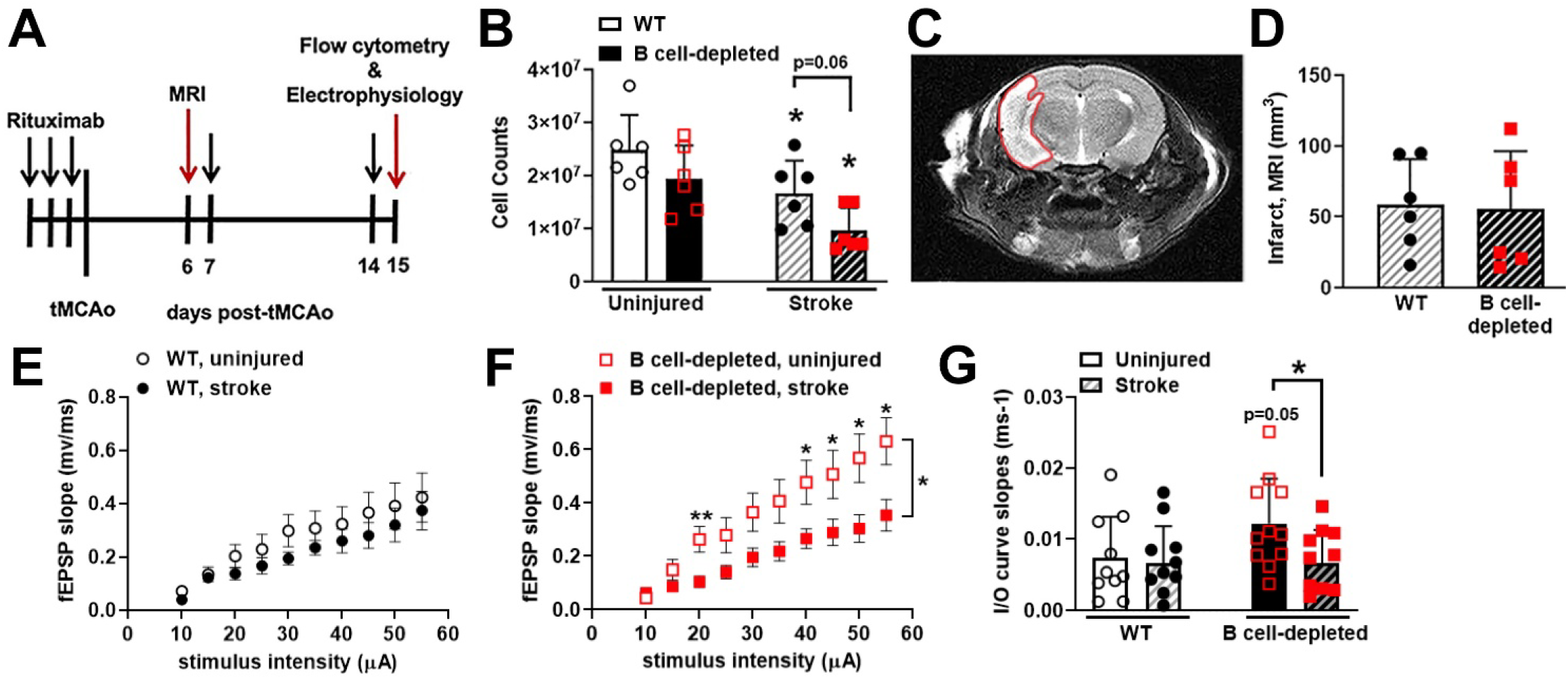
Stroke reduces synaptic transmission in B cell-depleted mice. (A) Experimental 125 timeline for electrophysiology recordings in the dentate gyrus (DG). Uninjured cohorts are solid bar graphs (WT, white; B cell depleted, black) and post-stroke cohorts have hatched bar graphs. (B) Stroke reduced the number of splenic CD19+ B cells in WT (hCD20-; black closed circles) mice compared to uninjured WT mice (black open circles), with a reduction in post-stroke B 129 cell-depleted mice (hCD20+; red closed squares) compared to uninjured B cell-depleted mice 130 (open red squares). (C) Representative MRI image identifying hyper-intense areas (with high 131 water content due to dilated ventricles and edema) outlined in red 6 days after stroke. (D) MRI infarct volume quantification of WT and B cell-depleted mice. (E-G) Electrophysiology generated input-output curves measuring transmission at perforant path-DG synapses in (E) WT and (F) B 134 cell-depleted mice, showing an effect of B cell depletion when (G) comparing individual I-O 135 curve slopes (each data point represents slope of the I-O curve from one brain slice). 136 Significance determined by (D) Student’s t-test, (E, F) repeated measures two-way ANOVA or (B, G) (*p<0.05, **p<0.01 vs. uninjured or WT control unless indicated by bracket), n=6 mice per 138 group, 2-4 brain slices per mouse.

### Establishing wide-field calcium imaging to investigate the modulation of neuronal function by circulating B cells

We next used both male and female mice of various ages (range of 2-20 months old) within an extended post-stroke timepoint of 3 weeks to examine sex differences in the context of the known age-dependent decrease in post-stroke neurogenesis.^17,18^ Further, the 3 week timepoint matches the known maturation and integration timeline of new granule cells receiving excitatory inputs and connecting to CA3 pyramidal cells.^19^ A total of 12,065 cells were individually recorded (6,875 from the DG; 5,190 from the CA1) from n=48 animals (26 males, 22 females; mean age 10.99 ± 5.38 months) and examples of GCaMP6s fluorescence are in **Figure 2A-C**. Further group breakdowns are found in **Supplemental Table 1** and shown by sex, B cell depletion status, infarct averages, animal counts, and analyzed cell counts. Neither age nor sex impacted the number of cells recorded in uninjured mice. In post-stroke animals, we verified that 1) systemic B cell depletion did not significantly alter the number of recorded cells, and 2) large infarcts with hippocampal involvement did not affect the number of recorded cells in the contralesional hippocampus. Finally, we tested for sex effects in the number of post-stroke cells recorded in the DG and CA1 and found no significant differences.

**Figure 2:**
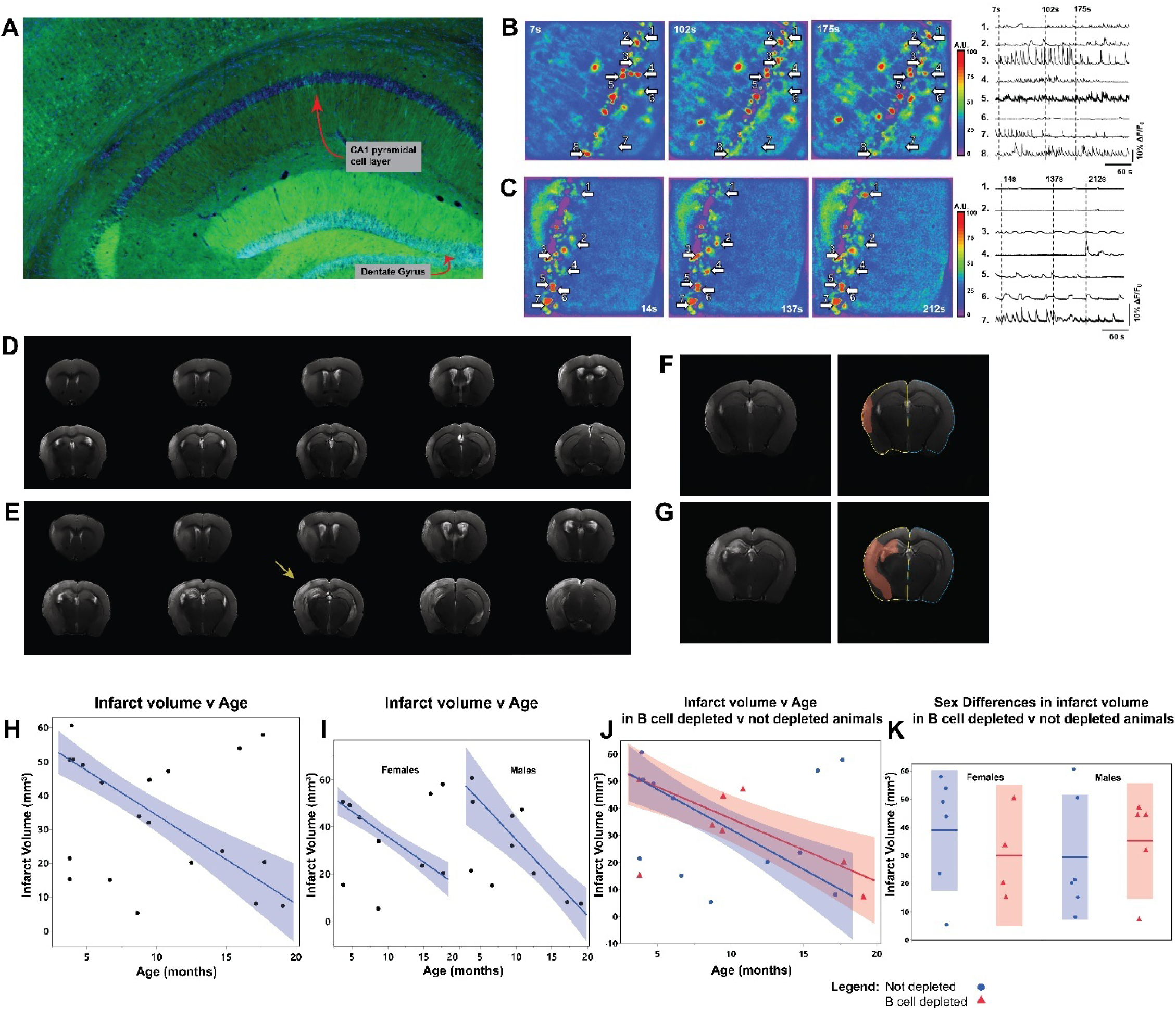
Representative figure of Ca^2+^ fluorescence, MRI, and infarct volumes. (A) Epifluorescence image of recording locations in a Synapsin-Cre x GCaMP6s mouse. Male mouse. DAPI in blue, with green showing GCaMP6s expression. (B-C) Pseudo-color heatmaps of neuronal calcium fluorescence within the same field of view at time points indicated at the bottom right of each panel. Top row shows CA1 (B), and bottom row shows DG (C). Areas of low activity are shown in cooler colors (blue) and areas of high acivity are depicted in warmer colors (red) representing the arbitrary units (A.U) of fluorescence displayed in the scale bar to the right. The units (A.U) on the scale bar represent fractions of maxium fluorescense intensity across the entire field of view (FOV). Right: Representative ΔF/F_0_ traces, displaying calcium activity across the imaging session (300 seconds) in select neurons indicated by the arrows. (D) 18-month-old female mouse (E) 9-month-old male mouse. Yellow arrow indicates large infarct with hippocampal involvement. (F and G) Representative examples of infarct volume analysis. Red indicates the infarct, yellow outline shows the ipsilesional hemisphere, blue shows the contralesional hemisphere. (H) Infarct volume v Age. (I) Sex differences on infarct volume v age. (J) Infarct volume v age in B cell depleted animals. (K) Sex differences in infarct volume in B cell depleted v not depleted animals. H, I, and J used a robust regression with the Robust Cauchy method in JMP Pro. This approach applies a weighting scheme that diminishes the impact of observations with large residuals, providing more reliable parameter estimates. (H) ChiSq(1) 37.71, p<0.0001. (I) Females ChiSq(1) 50.75, p<0.0001; Males ChiSq(1) 32.44, p<0.0001. (J) Not Depleted ChiSq(1) 20.38, p<0.0001; B cell depleted ChiSq(1) 19.66, p<0.0001. No significant effect of B cell depletion on infarct volume. (K) No sex effects on infarct volumes in animals.

However, taking age at stroke onset into account, older post-stroke mice had fewer cells recorded in the contralesional DG (β=-3.74, p=0.004) but with no effect in the CA1 (β=-1.56, p=0.32).

We evaluated infarct volumes using MRI performed at one-week post-stroke to determine if lesion size influenced calcium activity (**Figures 2D-G**). Overall, mean infarct volume was unaffected by sex and B cell depletion (**Figures 2H-K**), with only advancing age correlating with smaller infarct sizes after tMCAo, corroborating existing literature.^20–24^ It should be noted that all infarct volumes over 31.987 mm^3^ had hippocampal involvement and occurred in younger mice. When analyzing median values of amplitude and frequency of calcium events, ipsilesional hippocampal involvement in the area of ischemic injury also did not have a significant effect in either DG or CA1 imaging data.

### Systemic B cell depletion alters neuronal calcium activity in the DG of unijured mice

For this data set, we recorded activity from 6,875 DG cells, with 908 cells from the DG in uninjured control animals (n=14 mice) and 3,148 DG cells from B cell-depleted uninjured animals (n=11 mice). Overall statistical results are shown in **Supplement Table 2**, with B cell depletion inducing significant main effects on the amplitude of calcium transients (p<0.0059) and interacting with Age (p<0.0036) and Sex x Age (p<0.038). B cell depleted groups showed a decreased trend in amplitudes in median values (p=0.0507; **Figure 3A**) and individual cell values (p<0.0001; **Figure 3C**) in the DG. Determining sex differences, only neurons from B cell-depleted females exhibited lower amplitudes for both median (p=0.0321) and cell values (p<0.0001; **Figure 4A,B**). We also observed that B cell depletion increased the frequency of calcium events in uninjured mice as both B cell-depleted females and males had higher event frequency compared to their non-depleted counterparts (p<0.0001, and p<0.01, respectively; **Figures 4C,D**). Notably, B cell depletion affected both males and females, though the decrease in calcium amplitudes were more robust in females, highlighting a sex-influenced regulation of DG activity by depletion of circulating B cells.

**Figure 3:**
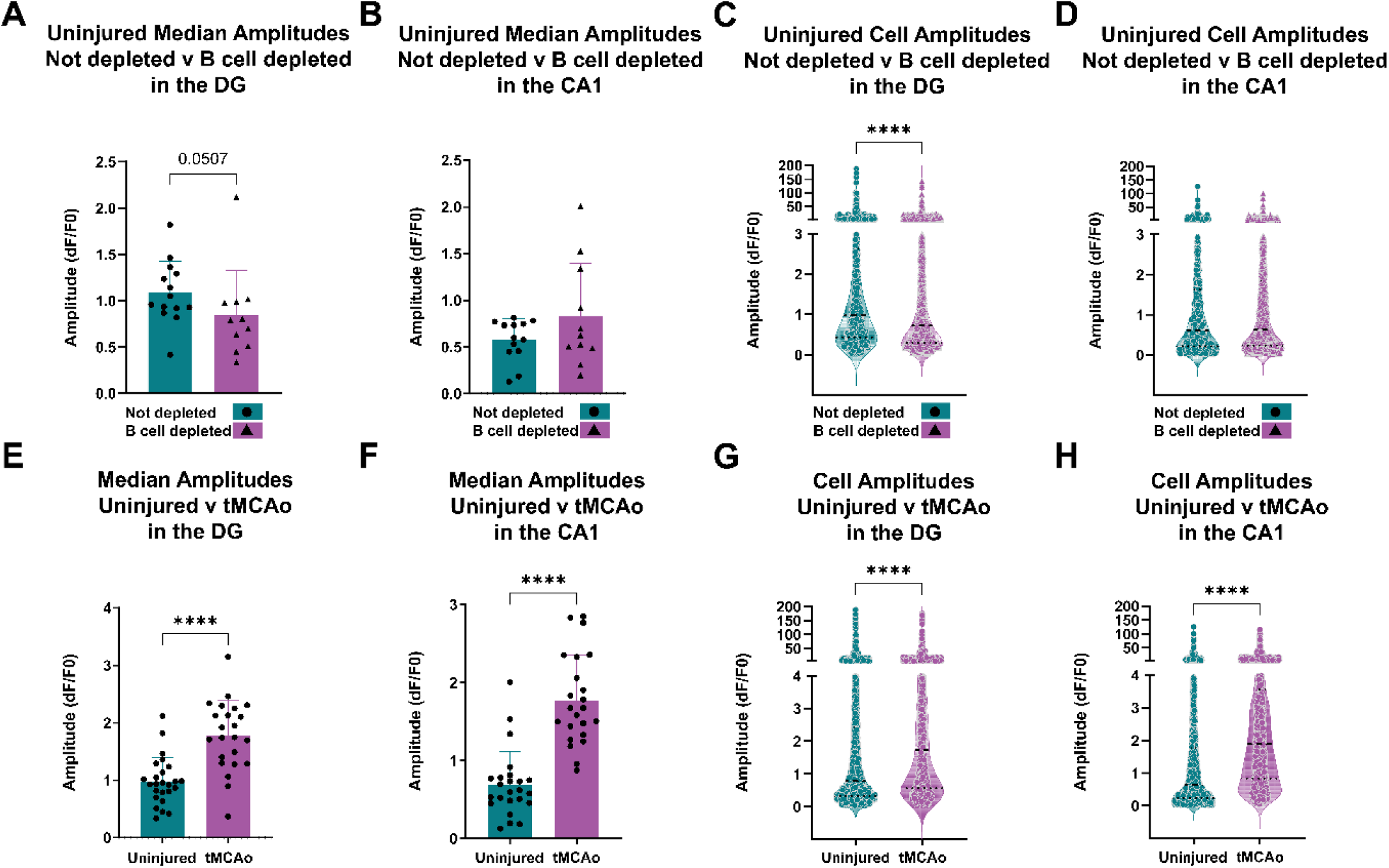
Effect of B cell depletion on neuronal activity in the DG and CA1. (A) Median amplitude values per animal in the DG. (B) Median amplitude values per animal in the CA1. (C) All cell amplitudes in the DG. (D) All cell amplitudes in the CA1 (E) Median amplitude values per animal in the DG (F) Median amplitude values per animal in the CA1 (G) All cell amplitudes in the DG (H) All cell amplitudes in the CA1. *p<0.05, **p<0.01, ***p<0.001, ****p<0.0001 as determined using a Mann-Whitney test for nonparametric data. For A,B,E,F bar graph presents means with SD. For C,D,G,H violin plots show all data points, with the main dotted line representing median. (A) Mann-Whitney, p=0.0507. (B) Not significant. (C) Mann-Whitney, p < 0.0001. (D) Not significant. (E-F) Mann-Whitney, p < 0.0001.

**Figure 4:**
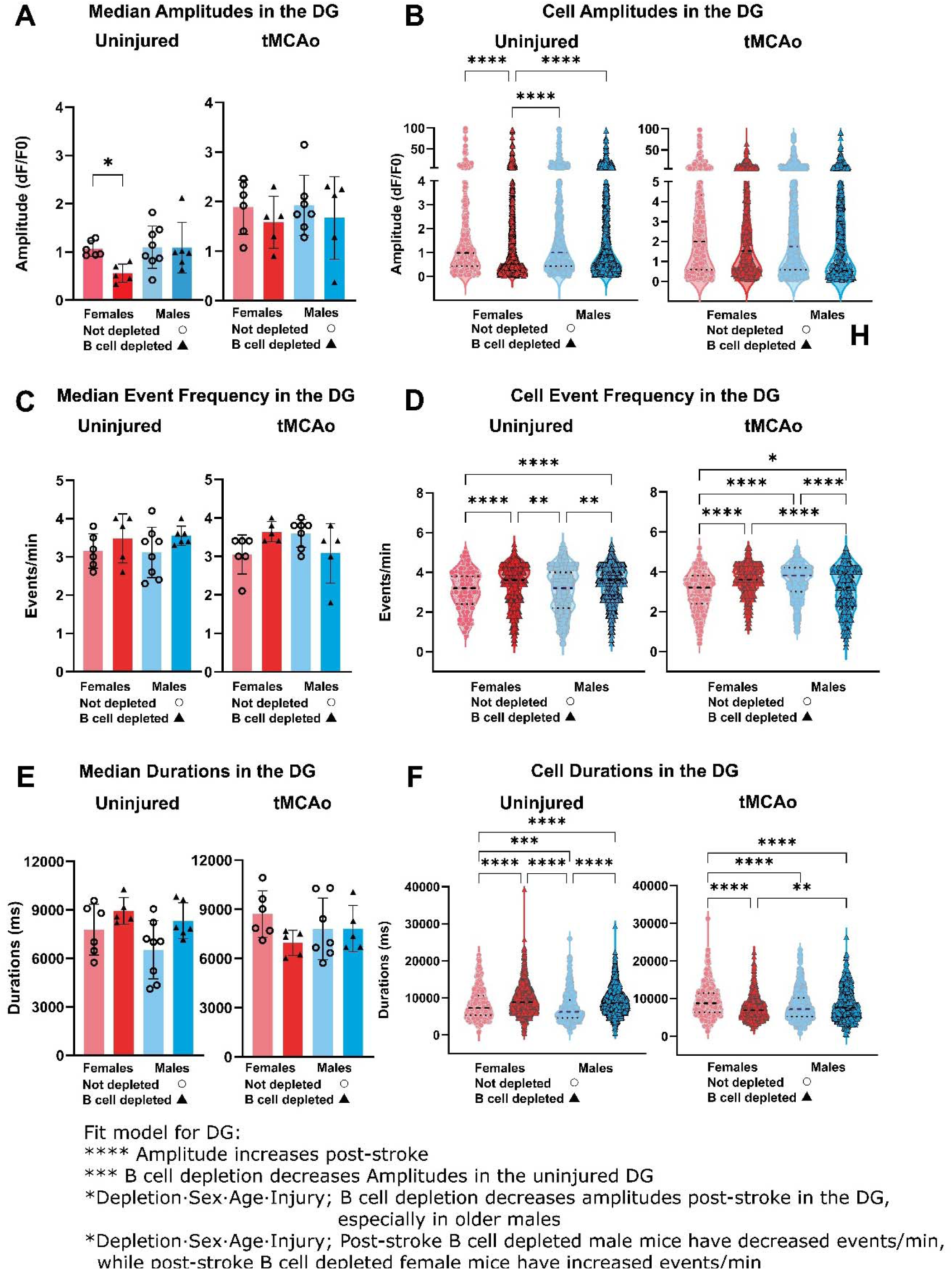
Sex differences in amplitudes, event frequency, and durations in the contralesional dentate gyrus. The graphs on the left are for uninjured, right are post-stroke (tMCAo). (A) Median amplitude values per animal in the DG, in uninjured and tMCAo. (B) All cell amplitudes in the DG, in uninjured and tMCAo. (C) Median events per minute per animal in the DG, in uninjured and tMCAo. (D) All cell events per minute in the DG, in uninjured and tMCAo. (E) Median durations in the DG, in uninjured and tMCAo. (F) All cell durations in the DG, in uninjured and tMCAo. For (A,C,E) bar graph presents means with SD. For (B,D,F) violin plots show all data points, with the main dotted line representing median. *p<0.05, **p<0.01, ***p<0.001, ****p<0.0001 as determined using a Kruskal-Wallis test for nonparametric data, along with Dunn’s multiple comparisons test. (A) Uninjured, Kruskal-Wallis test, p=0.0321, with multiple comparisons shown on graph. tMCAo, not significant (B) Uninjured, Kruskal-Wallis test, p<0.0001, with multiple comparisons shown on the graph. tMCAo, not significant. (C) Not significant. (D) Kruskal-Wallis test, p<0.0001, for both uninjured and tMCAO, with multiple comparisons shown on the graph. (E) Not significant. For uninjured, Kruskal-Wallis test, p=0.0657. (F) Kruskal-Wallis test, p<0.0001, for both uninjured and tMCAO, with multiple comparisons shown on the graph. At the bottom of the figure all fit model stats are listed, with further statistical tables in supplement.

### Post-stroke B cell depletion reduces Ca^2+^ event amplitudes in the DG and differentially alters event frequency in male versus female mice

As mentioned previously, all stroke mice (n=23) underwent a 45-min tMCAo 3 weeks prior to calcium imaging. A total of 2,819 cells were recorded in the contralesional DG post-stroke and, when compared to DG cells recorded from uninjured mice, stroke produced a robust increase in contralesional DG calcium amplitudes (p<0.0001; **Figure 3E,G**). Overall statistical interactions confirm a main effect of stroke injury increasing median DG amplitudes (p<0.0001; **Supplemental Table 2**), with complex interactions between Sex x Age (p=0.0001), Depletion x Age (p=0.0417) and a four-way interaction with Depletion, Sex, Age, and Injury (p=0.0107). Post-stroke B cell depletion lowered amplitudes in the DG for both females and males (**Figure 4A,B – right graphs**) to values closer to non-injured mice. Post-stroke B cell-depleted male mice also exhibited lower DG amplitudes with increasing age (**Figure S4A**). In contrast, B cell depletion in post-stroke female mice led to higher calcium events frequencies (p<0.0001; **Figure 4C,D**) whereas event frequency was decreased in male post-stroke B cell depleted mice, especially with increasing age (p=0.0308) when accounting for age in our fit model (Depletion x Sex x Age x Injury). These data provide further support for a sex-dependent dichotomy for changes in DG activity in response to injury and B cell depletion.

### Examining unique trends in DG calcium activity

To aid interpretation of calcium signals, we assessed event duration alongside amplitude and frequency. In Figure 4A, uninjured female mice with B cell depletion exhibited the lowest calcium event amplitudes. This pattern did not extend to event frequency. Instead, B cell depleted uninjured females showed higher median event frequencies than their non-B cell depleted counterparts (**Figure 4C,D**). This group also exhibited increased transient duration based on both median and per-cell measures (**Figure 4E,F**). We calculated calcium activity index (transient amplitude x duration x event frequency) as well as calcium load (amplitude x duration)^16^ to aid in understanding these trends (**Supplement Figure 3**) and confirm the most robust change in calcium load with B cell depletion in uninjured females. Additionally, we examined age effects for DG amplitudes and noted that both sexes trend upwards with age in non-B cell depleted conditions, that is lost upon B cell depletion (**Supplementary Figure 4A**), including that B cell depleted males post-stroke exhibited a decrease in calcium amplitudes.

### Age Effects on Hippocampal Calcium Transients

Stroke outcomes and neuroinflammatory trajectories are age-dependent, and adaptive immune compartments shift with aging in ways that could plausibly alter hippocampal activity; therefore, we incorporated age as a key biological variable in the study.^23,24^ Aging is associated with increased baseline inflammatory tone and heightened immune cell reactivity, while B-cell populations skew toward more antigen-experienced and pro-inflammatory phenotypes.^25,26^ These have been linked to worse neurological recovery and altered synaptic plasticity after ischemic injury.^27,28^ In the DG, median calcium transient amplitudes in non–B cell depleted mice tended to increase with age in both sexes, whereas this relationship was absent following B-cell depletion (**Supplementary Figure 4A**). In fact, post-stroke B cell depleted males and females exhibited decreased amplitudes with increasing age. Collectively, these patterns are consistent with the interpretation that circulating B cells contribute to age-related, sex-dependent modulation of hippocampal calcium signaling after stroke.

### B cell depletion increases CA1 calcium event amplitudes in uninjured males but reduces event frequency across all cell groups

One of the benefits of using *ex vivo* slice imaging was the ability to also quantify within-animal changes to CA1, a hippocampal region that, unlike the DG, did not experience significant B cell diapedesis at 4 days after stroke.^1^ For the CA1, we recorded activity from 5,190 total cells. These were split by 2,003 cells from uninjured control mice (n=25 mice; 11 female, 14 male) and 3,187 cells in the post-stroke groups (n=23 mice, 11 female, 12 male). Unlike in the DG, grouped data showed no differences in calcium event amplitudes with B cell depletion (**Figure 3B,D**). Overall statistical interactions, however, did demonstrate a main effect of Sex on CA1 calcium event amplitude (p=0.0074; **Supplemental Table 3**), with trending effects for Depletion (p=0.0598) on amplitude as B cell-depleted males showed higher amplitudes than all other groups (**Figure 5A,B**). B cell depletion lowered event frequency in the CA1 exhibiting a trend for a Depletion x Sex effect (p=0.0558) with reduced event frequency upon B cell depletion greatest in female mice (**Figure 5C,D**).

**Figure 5:**
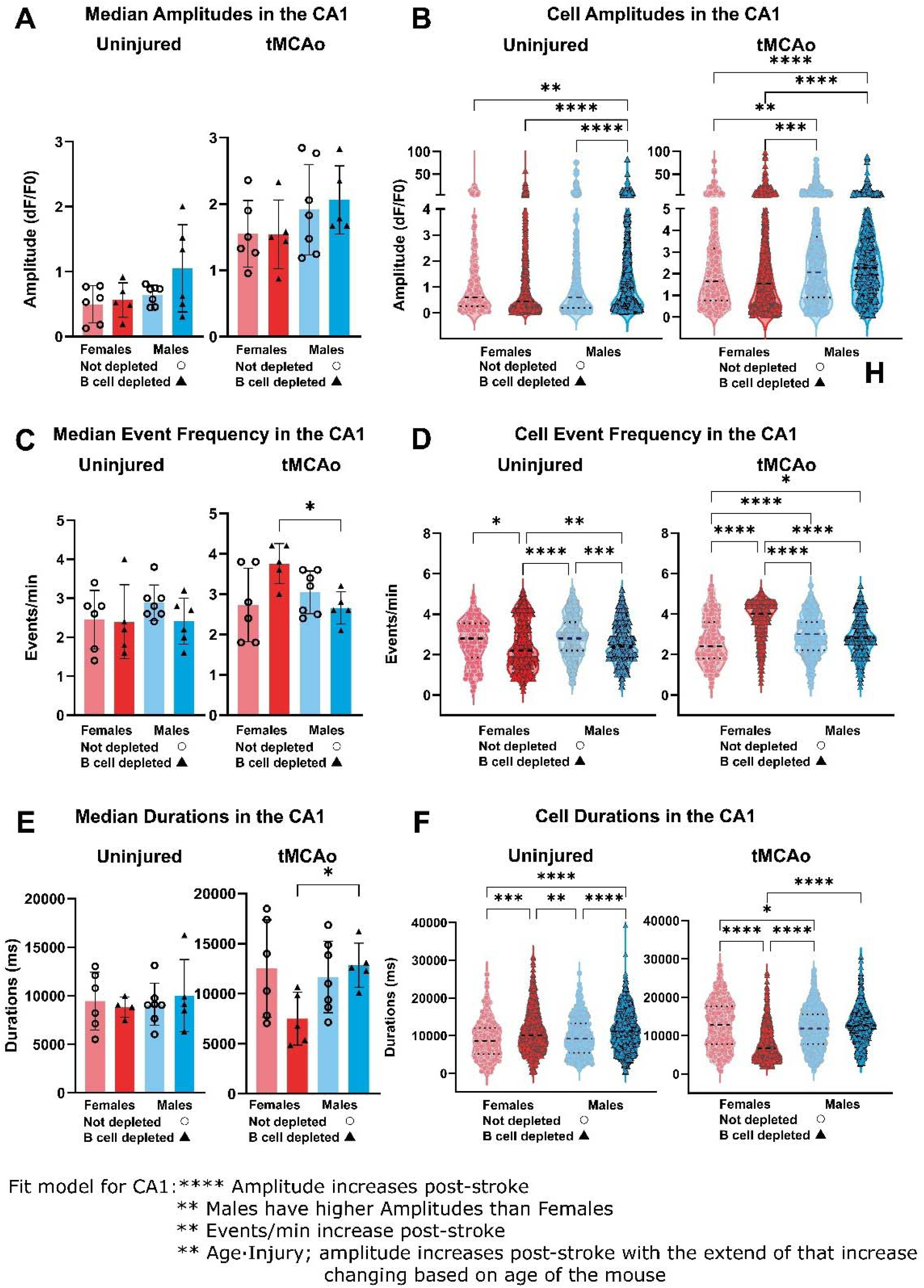
Sex differences in amplitudes, event frequency, and durations in the contralesional CA1. The graphs on the left are for uninjured, right are post-stroke (tMCAo). (A) Median amplitude values per animal in the CA1, in uninjured and tMCAo. (B) All cell amplitudes in the CA1, in uninjured and tMCAo. (C) Median events per minute per animal in the CA1, in uninjured and tMCAo. (D) All cell events per minute in the CA1, in uninjured and tMCAo. (E) Median durations in the CA1, in uninjured and tMCAo. (F) All cell durations in the CA1, in uninjured and tMCAo. For (A,C,E) bar graph presents means with SD. For (B,D,F) violin plots show all data points, with the main dotted line representing median. *p<0.05, **p<0.01, ***p<0.001, ****p<0.0001 as determined using a Kruskal-Wallis test for nonparametric data, along with Dunn’s multiple comparisons test. (A) Not significant (B) Kruskal-Wallis test, p<0.0001, for both uninjured and tMCAO, with multiple comparisons shown on the graph. (C) Uninjured Not significant. For tMCAo, Kruskal-Wallis test, p=0.0473, multiple comparisons shown on the graph. (D) Kruskal-Wallis test, p<0.0001, for both uninjured and tMCAO, with multiple comparisons shown on the graph. (E) For uninjured, not significant. For tMCAo, p=0.0423, with multiple comparisons shown on the graph (F) Kruskal-Wallis test, p<0.0001, for both uninjured and tMCAO, with multiple comparisons shown on the graph. At the bottom of the figure all fit model stats are listed, with further statistical tables in supplement.

### Stroke injury increases CA1 neuronal amplitudes and event frequency

We recorded from 3,187 cells specifically in the post-stroke CA1 (1,597 from males, 1,590 from females) and identified a stronger increase in contralesional hippocampal CA1 amplitudes after stroke for grouped data (both p<0.0001; **Figure 3F,H**) versus the DG. This was also confirmed in the multiple linear regression model, with a main effect of stroke (p<0.0001; **Supplement Table 3**) and Age x Injury (p=0.0034) on CA1 neuronal amplitudes. In cell CA1 values, males exhibited elevated amplitudes compared to their female counterparts (p<0.0001; **Figure 5A,B**). There were no post-stroke interactions with B cell depletion as found in the DG, but as expected given the lack of robust B cell recruitment into this region in our prior work.^1^ Also unlike the DG, which had no change in post-stroke event frequency, the CA1 of pooled post-stroke mice exhibited higher median (p=0.0235) and cell (p<0.0001) firing rates at 3 weeks after tMCAO. For the event frequency, there was a significant interaction in median values in the CA1 (p=0.0275; **Figure 5C**) as well as in cell numbers (p<0.0001; **Figure 5D**) as female B cell-depleted animals exhibited the highest CA1 firing events/min after stroke.

### Examining unique trends in CA1 calcium activity

In CA1, B cell depleted post-stroke female mice exhibited a distinct pattern. This group showed the highest event frequency (**Figure 5C,D**) while also displaying the shortest event durations (**Figure 5E**,**F**) across all groups, including uninjured controls. When these measures were integrated into composite metrics (**Supplemental Figure 3B,D**), the opposing effects offset one another, yielding no net difference. Additionally, we examined the effects of aging on calcium amplitudes. We show that post-stroke CA1 amplitudes tend to decline with age, though this effect was not as robust with B cell depletion (**Supplemental Figure 4B**). Post-stroke B cell depleted males decreased less with age than their non-depleted counterparts, while post-stroke females also tend to show decreased amplitudes with increasing age.

## Discussion

Overall, our study shows that in the tMCAo model of ischemic stroke, injury, age, sex, and B cell depletion interact in complex ways to affect neuronal function in multiple hippocampal regions (data summarized in **Figure 6**). Perhaps the most surprising finding of this study is that systemic B cell depletion, in the absence of any injury and regardless of age or sex, significantly alters neuronal activity in the hippocampal DG. This is a novel observation that challenges the traditional view of B cells as solely ‘peripheral immune mediators’. It suggests that B cells, even in a healthy state, play a tonic neuromodulatory role in brain regions critical for cognition and memory.

**Figure 6.**
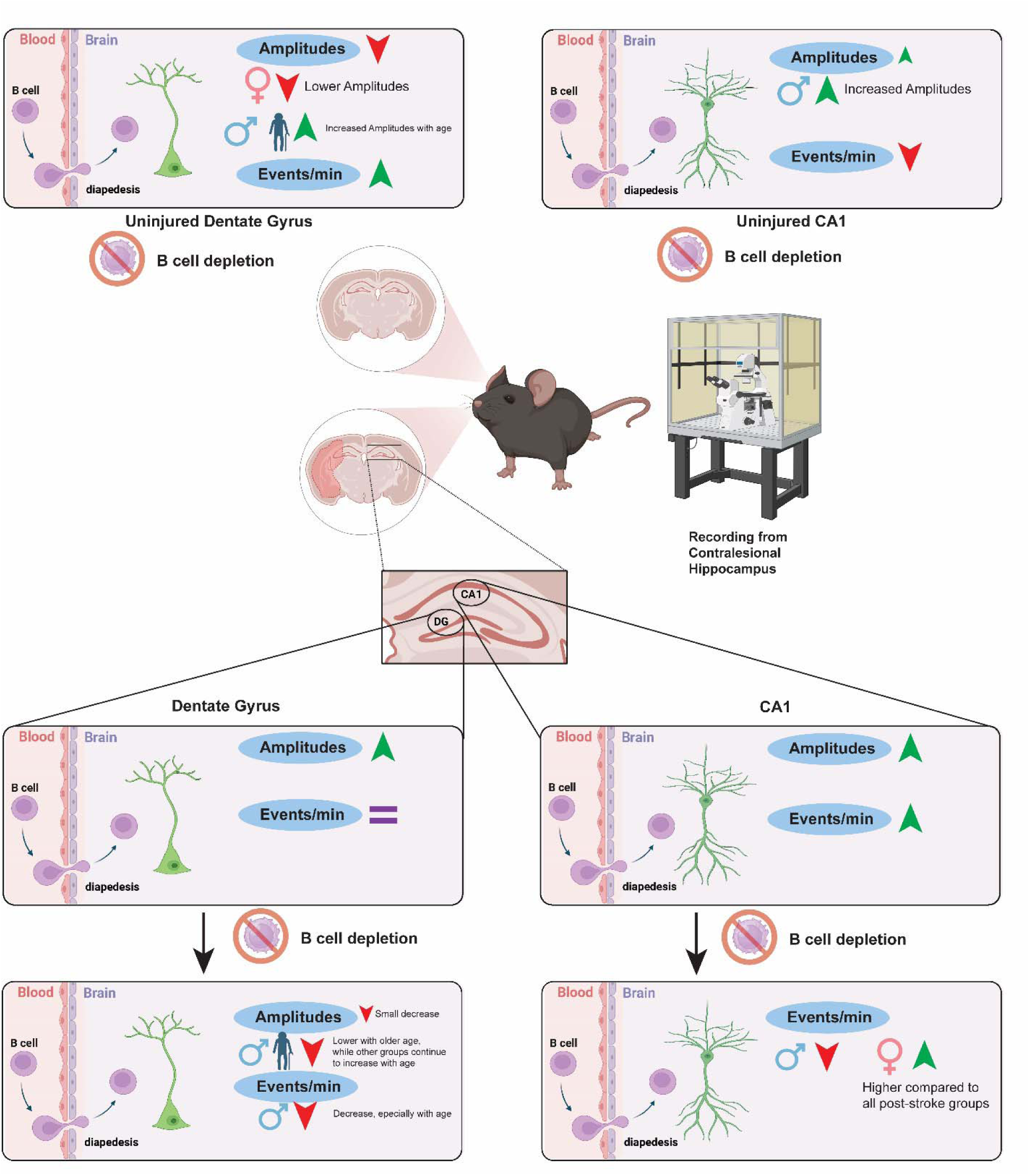
Summary of results.

B cells can be neurotrophic, as they are the primary lymphocytic source of brain-derived neurotrophic factor (BDNF), which supports neuronal function and plasticity.^29^ Our prior work found that brain-localized B cells sense glutamate and secrete mature BDNF, promoting neuronal survival and limiting brain injury after stroke, including in aged mice.^30^ In addition to being a localized source of BDNF production, B cells may alter hippocampal signaling through cytokine production and/or modulation of glial cells^31^ as they produce cytokines including IL-10, IL-6, TNF-α, TGF-β, which modulate neuronal signaling,^31^ specifically with IL-10 and TGF-β inducing neuroprotective effects and supporting hippocampal long-term potentiation.^32^

We also observed that ischemic injury had significant effects on contralesional neuronal activity, with B cell depletion modulating DG dynamics in sex-, age-, and injury-dependent manners. Following ischemic stroke, there is acute neuronal death and neuroinflammation, followed by cell proliferation that facilitates tissue replacement and remodeling.^33^ Among brain regions affected, the hippocampus is especially vulnerable after stroke.^34–36^ Quite often, post-stroke impairment is associated with hippocampal neurodegeneration and decreased hippocampal function,^37^ with male mice exposed to a 45-min tMCAo exhibiting hippocampal LTP impairment through at least 30 days after stroke.^36^ Furthermore, reduced LTP was both ipsilesional and contralesional, with no statistical difference between the two.^36^ In our study, we observed that tMCAo alone significantly increased calcium transient amplitudes in both the contralesional CA1 and DG regions. Interestingly, CA1 pyramidal neurons in the contralesional hippocampus of tMCAo-treated mice exhibit higher neuronal complexity and lower density of spines,^4^ with spines being where most excitatory synapses are located.^38^ Thus, a reduction in spine density implies that the neuron is receiving fewer excitatory inputs^39^ and in response, neurons may engage in compensatory mechanisms to preserve normal firing rates.^40^ One form of intrinsic plasticity involves lowering the firing threshold, thereby increasing the cell’s sensitivity to remaining excitatory inputs.^41^ Thus, the post-tMCAo increases in calcium transient amplitude in CA1 and DG, together with increased CA1 event frequency, may indicate compensatory circuit hyperexcitability following stroke, arising from reduced afferent input from the injured hemisphere and/or neuronal loss within vulnerable hippocampal subpopulations. This may not, however, be a sign of healthy recovery but instead disorganized firing that could ultimately contribute to long-term cognitive deficits and seizure susceptibility.^2,42^

Prior work highlights that B cells can exert time- and context-dependent effects after stroke: they may be protective early, yet later contribute to post-stroke cognitive impairment through immune mechanisms such as cytokine production, antibody secretion, and modulation of T cell responses.^43^ Consistent with this non-uniform role, our data show that circulating B cells differentially shape hippocampal network activity in a manner that depends on sex, injury status, and age, with the strongest effects of B cell depletion emerging in aged mice after stroke. This age sensitivity is notable given evidence that B cells can be linked to loss of hippocampal synaptic plasticity and cognitive decline, potentially via autoimmune responses to CNS antigens that may be amplified in pro-inflammatory states.^44^ Aged mice have worse behavioral impairments 2 weeks post-stroke,^24^ and aged post-stroke male mice experience respiratory and cognitive deficits.^45^ Aging is associated with slower maturation and poor circuit integration of newborn neurons, which could also lead to more aberrant neurogenesis post-stroke and worse cognitive outcomes.^46^ Our findings suggest that the impact of depleting circulating B cells is likely shaped by the aged post-stroke immune milieu, where post-stroke immunosuppression combined with immunosenescence could shift the balance toward maladaptive consequences^47^ (increased infection susceptibility, and/or impaired cognitive recovery), potentially through mechanisms that are not limited to antibody production.

One key thing to note is hippocampal regional differences in the effect of B cells on neuronal function. The DG appeared more sensitive to modulation by B cell depletion and age-related effects versus the CA1 region. This may reflect underlying heterogeneity between hippocampal subregions, including differences in inhibitory tone, constitutive neurogenesis, or immune accessibility. Other studies have also shown that DG is one of the subregions that is most vulnerable to aging,^48,49^ possibly due to the reduced synaptic contacts from the entorhinal cortex (EC).^50^ Also, a sharp decline in neurogenesis with aging could be a key factor as well in regional neuronal function.^51,52^ As mentioned, our prior study showed significant B cell diapedesis into the contralesional DG at 4 days after tMCAo in young male mice.^1^ By depleting B cells systemically, we are likely seeing a stronger effect in the DG due to the possibility that the region would have more B cells post-stroke than the CA1, since only the DG exhibited significant diapedesis in our prior study,^1^ but in this work we did not quantify the number of B cells in comparison between CA1 and DG which should be a focus of future studies.

We observed that age increased calcium transient amplitudes in both CA1 and DG, but only when interactions involved injury. However, in B cell depleted post-stroke mice, DG calcium transient amplitude decreased with age, especially in older males. This was not evident in the CA1, again highlighting regional differences. A calcium imaging study of young (3-4mo.) and old (21-26mo.) female mice found that calcium activity was significantly higher in the DG of the older mice and that the aged DG was hyperexcitable.^53^ This was similar to what we observed in post-stroke mice without B cell depletion. This hyperexcitable DG with aging can possibly reflect a compensation mechanism for reduced EC inputs.^53^ Also, this age related hyperexcitability may be a maladaptive homeostatic mechanism that is exacerbated by stroke inflammation and sensitive to peripheral B cell depletion. Studies show that the loss of afferent inputs to cortical neurons leads to enhanced excitatory synaptic strength and increased intrinsic excitability.^54^ In a mouse study of penetrating brain injury, there is compensation for lost neurons that leads to an increase in excitation.^55^ Furthermore, in the context of stroke, homeostatic plasticity results in hyperexcitability over the first week to one month after recovery,^56^ while in a mouse model of ischemic stroke, the contralesional hemisphere also showed gradual hyperexcitability in the somatosensory cortex after contralateral stimulation.^57^ As mentioned, future work needs to carefully distinguish which hippocampal adaptations (neurogenesis, changes in synaptic transmission, cellular or network excitability) are directly linked to post-stroke cognition, and how the systemic immune response can modulate these effects.

In summary, our mouse model identifies circulating B cells as modulators of hippocampal network activity, with effects that depend on sex, age, and injury status. These data advance a view of immune cells, like B cells, as not only responders to CNS injury, but active players in tuning neural excitability and plasticity. In fact, B cells are central players to functional recovery whose role has been evolving and shifting over time, from beneficial and neurotrophic to maladaptive and detrimental depending on timing, context, subset, and immune state. These findings add, however, to a more unified model of neuro-immune interactions that opens the door for novel immunotherapies that can harness supportive functions of B cells while slowing their maladaptive roles in stroke and aging.

## Acknowledgments

These studies are supported by grants from the Dana Foundation David Mahoney Neuroimaging Program and the National Institutes of Health to AMS: R01NS088555, RF1NS088555; to TU: T32NS077889, 3RF1NS088555-07A1S1; to VOT: 3R01NS088555-02S1, 5T32AI005284-40; to NM: NS102417; to PO: R01DA041513.

## References

1 Ortega, S. B. et al. B cells migrate into remote brain areas and support neurogenesis and functional recovery after focal stroke in mice. Proc Natl Acad Sci U S A 117, 4983–4993 (2020). 10.1073/pnas.1913292117

2 Cuartero, M. I. et al. Abolition of aberrant neurogenesis ameliorates cognitive impairment after stroke in mice. J Clin Invest 129, 1536–1550 (2019). 10.1172/JCI120412

3 Wang, C. et al. Sustained increase in adult neurogenesis in the rat hippocampal dentate gyrus after transient brain ischemia. Neurosci Lett 488, 70–75 (2011). 10.1016/j.neulet.2010.10.079

4 Merino-Serrais, P. et al. Structural changes of CA1 pyramidal neurons after stroke in the contralesional hippocampus. Brain Pathol 34, e13222 (2024). 10.1111/bpa.13222

5 Lee, C. H. et al. Long-term changes in neuronal degeneration and microglial activation in the hippocampal CA1 region after experimental transient cerebral ischemic damage. Brain Res 1342, 138–149 (2010). 10.1016/j.brainres.2010.04.046

6 Gemmell, E. et al. Hippocampal neuronal atrophy and cognitive function in delayed poststroke and aging-related dementias. Stroke 43, 808–814 (2012). 10.1161/STROKEAHA.111.636498

7 Werden, E. et al. Structural MRI markers of brain aging early after ischemic stroke. Neurology 89, 116–124 (2017). 10.1212/WNL.0000000000004086

8 Jung, J. et al. Altered hippocampal functional connectivity patterns in patients with cognitive impairments following ischaemic stroke: A resting-state fMRI study. Neuroimage Clin 32, 102742 (2021). 10.1016/j.nicl.2021.102742

9 Adhikari, Y., Ma, C. G., Chai, Z. & Jin, X. Preventing development of post-stroke hyperexcitability by optogenetic or pharmacological stimulation of cortical excitatory activity. Neurobiol Dis 184, 106233 (2023). 10.1016/j.nbd.2023.106233

10 Monson, N. L. et al. Rituximab therapy reduces organ-specific T cell responses and ameliorates experimental autoimmune encephalomyelitis. PLoS ONE 6, e17103 (2011). 10.1371/journal.pone.0017103

11 Schneider, C. A., Rasband, W. S. & Eliceiri, K. W. NIH Image to ImageJ: 25 years of image analysis. Nat Methods 9, 671–675 (2012). 10.1038/nmeth.2089

12 Park, H. J. et al. Semi-automated method for estimating lesion volumes. J Neurosci Methods 213, 76–83 (2013). 10.1016/j.jneumeth.2012.12.010

13 Weber, R. Z. et al. A toolkit for stroke infarct volume estimation in rodents. Neuroimage 287, 120518 (2024). 10.1016/j.neuroimage.2024.120518

14 Ting, J. T. et al. Preparation of Acute Brain Slices Using an Optimized N-Methyl-D-glucamine Protective Recovery Method. J Vis Exp (2018). 10.3791/53825

15 Neugornet, A., O’Donovan, B. & Ortinski, P. I. Comparative Effects of Event Detection Methods on the Analysis and Interpretation of Ca(2+) Imaging Data. Front Neurosci 15, 620869 (2021). 10.3389/fnins.2021.620869

16 Patel, T. P., Man, K., Firestein, B. L. & Meaney, D. F. Automated quantification of neuronal networks and single-cell calcium dynamics using calcium imaging. J Neurosci Methods 243, 26–38 (2015). 10.1016/j.jneumeth.2015.01.020

17 Balseanu, A. T. et al. Electric Stimulation of Neurogenesis Improves Behavioral Recovery After Focal Ischemia in Aged Rats. Front Neurosci 14, 732 (2020). 10.3389/fnins.2020.00732

18 Yan, J., Liu, Y., Zheng, F., Lv, D. & Jin, D. Environmental enrichment enhanced neurogenesis and behavioral recovery after stroke in aged rats. Aging (Albany NY) 15, 9453–9463 (2023). 10.18632/aging.205010

19 Rodriguez-Iglesias, N., Sierra, A. & Valero, J. Rewiring of Memory Circuits: Connecting Adult Newborn Neurons With the Help of Microglia. Front Cell Dev Biol 7, 24 (2019). 10.3389/fcell.2019.00024

20 Wang, R. Y., Wang, P. S. & Yang, Y. R. Effect of age in rats following middle cerebral artery occlusion. Gerontology 49, 27–32 (2003). 10.1159/000066505

21 Liu, F., Yuan, R., Benashski, S. E. & McCullough, L. D. Changes in experimental stroke outcome across the life span. J Cereb Blood Flow Metab 29, 792–802 (2009). 10.1038/jcbfm.2009.5

22 Liu, F. & McCullough, L. D. Middle cerebral artery occlusion model in rodents: methods and potential pitfalls. J Biomed Biotechnol 2011, 464701 (2011). 10.1155/2011/464701

23 Ritzel, R. M. et al. Aging alters the immunological response to ischemic stroke. Acta Neuropathol 136, 89–110 (2018). 10.1007/s00401-018-1859-2

24 Manwani, B. et al. Functional recovery in aging mice after experimental stroke. Brain Behav Immun 25, 1689–1700 (2011). 10.1016/j.bbi.2011.06.015

25 Li, X. et al. Inflammation and aging: signaling pathways and intervention therapies. Signal Transduct Target Ther 8, 239 (2023). 10.1038/s41392-023-01502-8

26 de Mol, J., Kuiper, J., Tsiantoulas, D. & Foks, A. C. The Dynamics of B Cell Aging in Health and Disease. Front Immunol 12, 733566 (2021). 10.3389/fimmu.2021.733566

27 Wu, S. et al. Updates of the role of B-cells in ischemic stroke. Front Cell Neurosci 18, 1340756 (2024). 10.3389/fncel.2024.1340756

28 Malone, M. K. et al. The immunopathology of B lymphocytes during stroke-induced injury and repair. Semin Immunopathol 45, 315–327 (2023). 10.1007/s00281-022-00971-3

29 Vega, J. A., Garcia-Suarez, O., Hannestad, J., Perez-Perez, M. & Germana, A. Neurotrophins and the immune system. J Anat 203, 1–19 (2003). 10.1046/j.1469-7580.2003.00203.x

30 Torres, V. O. et al. B cells upregulate NMDARs, respond to extracellular glutamate, and express mature BDNF to protect the brain from ischemic injury. Neurobiol Dis 207, 106819 (2025). 10.1016/j.nbd.2025.106819

31 Aspden, J. W. et al. Intruders or protectors - the multifaceted role of B cells in CNS disorders. Front Cell Neurosci 17, 1329823 (2023). 10.3389/fncel.2023.1329823

32 Nenov, M. N., Malkov, A. E., Konakov, M. V. & Levin, S. G. Interleukin-10 and transforming growth factor-beta1 facilitate long-term potentiation in CA1 region of hippocampus. Biochem Biophys Res Commun 518, 486–491 (2019). 10.1016/j.bbrc.2019.08.072

33 Burda, J. E. & Sofroniew, M. V. Reactive gliosis and the multicellular response to CNS damage and disease. Neuron 81, 229–248 (2014). 10.1016/j.neuron.2013.12.034

34 Chapman, A. C. A place for place cells in post-stroke cognitive impairment. Trends Neurosci 48, 391–392 (2025). 10.1016/j.tins.2025.05.001

35 Heiser, H. et al. Brain-wide microstrokes affect the stability of memory circuits in the hippocampus. Nat Commun 16, 3462 (2025). 10.1038/s41467-025-58688-4

36 Orfila, J. E. et al. Delayed inhibition of tonic inhibition enhances functional recovery following experimental ischemic stroke. J Cereb Blood Flow Metab 39, 1005–1014 (2019). 10.1177/0271678X17750761

37 Gemmell, E. et al. Neuron volumes in hippocampal subfields in delayed poststroke and aging-related dementias. J Neuropathol Exp Neurol 73, 305–311 (2014). 10.1097/NEN.0000000000000054

38 Hering, H. & Sheng, M. Dendritic spines: structure, dynamics and regulation. Nat Rev Neurosci 2, 880–888 (2001). 10.1038/35104061

39 Amaral, M. D. & Pozzo-Miller, L. The dynamics of excitatory synapse formation on dendritic spines. Cellscience 5, 19–25 (2009).

40 Turrigiano, G. G. The self-tuning neuron: synaptic scaling of excitatory synapses. Cell 135, 422–435 (2008). 10.1016/j.cell.2008.10.008

41 Yang, J. & Prescott, S. A. Homeostatic regulation of neuronal function: importance of degeneracy and pleiotropy. Front Cell Neurosci 17, 1184563 (2023). 10.3389/fncel.2023.1184563

42 Cuartero, M. I. et al. Post-stroke Neurogenesis: Friend or Foe? Front Cell Dev Biol 9, 657846 (2021). 10.3389/fcell.2021.657846

43 Engler-Chiurazzi, E. B., Monaghan, K. L., Wan, E. C. K. & Ren, X. Role of B cells and the aging brain in stroke recovery and treatment. Geroscience 42, 1199–1216 (2020). 10.1007/s11357-020-00242-9

44 Doyle, K. P. et al. B-lymphocyte-mediated delayed cognitive impairment following stroke. J Neurosci 35, 2133–2145 (2015). 10.1523/JNEUROSCI.4098-14.2015

45 Patrizz, A. et al. Stroke-Induced Respiratory Dysfunction Is Associated With Cognitive Decline. Stroke 54, 1863–1874 (2023). 10.1161/STROKEAHA.122.041239

46 Rao, M. S., Hattiangady, B., Abdel-Rahman, A., Stanley, D. P. & Shetty, A. K. Newly born cells in the ageing dentate gyrus display normal migration, survival and neuronal fate choice but endure retarded early maturation. Eur J Neurosci 21, 464–476 (2005). 10.1111/j.1460-9568.2005.03853.x

47 Gallizioli, M., Arbaizar-Rovirosa, M., Brea, D. & Planas, A. M. Differences in the post-stroke innate immune response between young and old. Semin Immunopathol 45, 367–376 (2023). 10.1007/s00281-023-00990-8

48 Small, S. A., Chawla, M. K., Buonocore, M., Rapp, P. R. & Barnes, C. A. Imaging correlates of brain function in monkeys and rats isolates a hippocampal subregion differentially vulnerable to aging. Proc Natl Acad Sci U S A 101, 7181–7186 (2004). 10.1073/pnas.0400285101

49 Yassa, M. A., Mattfeld, A. T., Stark, S. M. & Stark, C. E. Age-related memory deficits linked to circuit-specific disruptions in the hippocampus. Proc Natl Acad Sci U S A 108, 8873–8878 (2011). 10.1073/pnas.1101567108

50 Burke, S. N. & Barnes, C. A. Senescent synapses and hippocampal circuit dynamics. Trends Neurosci 33, 153–161 (2010). 10.1016/j.tins.2009.12.003

51 Moreno-Jimenez, E. P. et al. Adult hippocampal neurogenesis is abundant in neurologically healthy subjects and drops sharply in patients with Alzheimer’s disease. Nat Med 25, 554–560 (2019). 10.1038/s41591-019-0375-9

52 Spalding, K. L. et al. Dynamics of hippocampal neurogenesis in adult humans. Cell 153, 1219–1227 (2013). 10.1016/j.cell.2013.05.002

53 McDermott, K. D., Frechou, M. A., Jordan, J. T., Martin, S. S. & Goncalves, J. T. Delayed formation of neural representations of space in aged mice. Aging Cell 22, e13924 (2023). 10.1111/acel.13924

54 Turrigiano, G. G., Leslie, K. R., Desai, N. S., Rutherford, L. C. & Nelson, S. B. Activity-dependent scaling of quantal amplitude in neocortical neurons. Nature 391, 892–896 (1998). 10.1038/36103

55 Ping, X. & Jin, X. Transition from Initial Hypoactivity to Hyperactivity in Cortical Layer V Pyramidal Neurons after Traumatic Brain Injury In Vivo. J Neurotrauma 33, 354–361 (2016). 10.1089/neu.2015.3913

56 Murphy, T. H. & Corbett, D. Plasticity during stroke recovery: from synapse to behaviour. Nat Rev Neurosci 10, 861–872 (2009). 10.1038/nrn2735

57 Barios, J. A. et al. Long-term dynamics of somatosensory activity in a stroke model of distal middle cerebral artery oclussion. J Cereb Blood Flow Metab 36, 606–620 (2016). 10.1177/0271678X15606139

