## Supplemental data for "Peripheral B cell populations tune spontaneous neuronal activity in the uninjured hippocampus after stroke"

Thomas A. Ujas^1^, Navid S. Tavakoli^1^, Pavel Yanev^2^, Vanessa O. Torres^3,4^, Jadwiga Turchan-Cholewo,^2^ Xiangmei Kong^3^, Erik J. Plautz^3^, Adam D. Bachstetter^1,5^, Nancy L. Monson^3^, Lenora J. Volk^6^, Pavel I. Ortinski^1^, and Ann M. Stowe^1,2,3,5^

^1^Department of Neuroscience, University of Kentucky, Lexington, Kentucky, USA

^2^ Department of Neurology, University of Kentucky, Lexington, Kentucky, USA

^3^Department of Neurology, University of Texas Southwestern Medical Center, Dallas, Texas, USA

^4^Denali Therapeutics Inc.; South San Francisco, California, USA 94080

^5^ Spinal Cord and Brain Injury Center, University of Kentucky, Lexington, Kentucky, USA

^6^ Department of Neuroscience, University of Texas Southwestern Medical Center, Dallas, Texas, USA

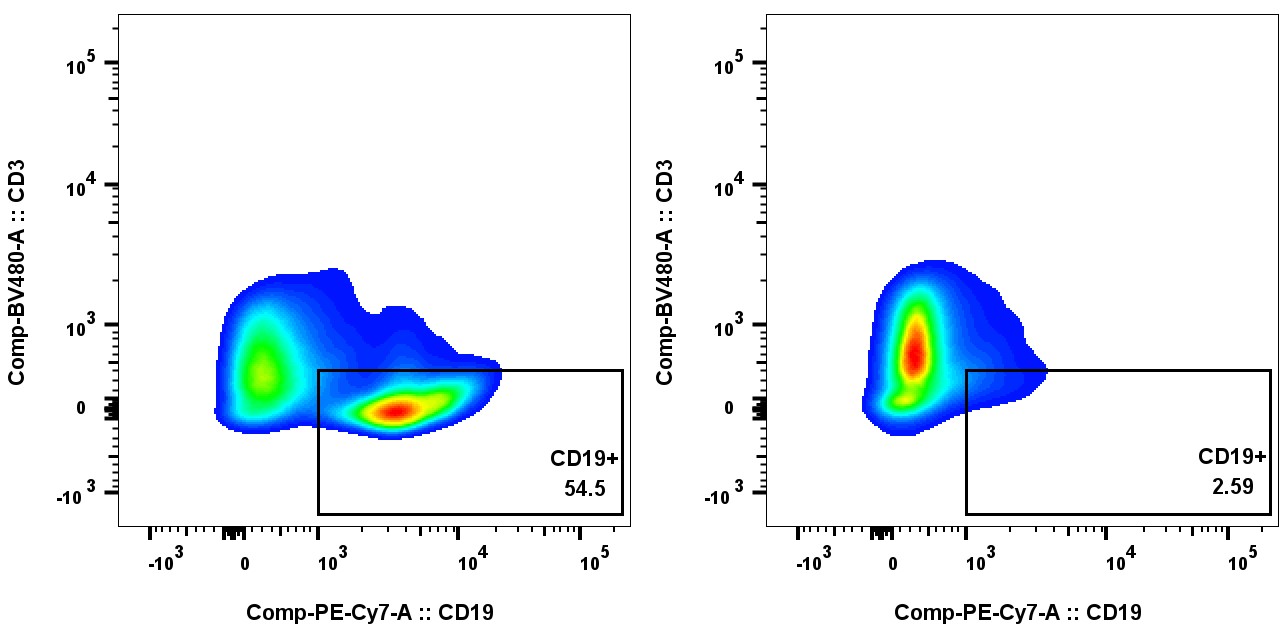

**Supplement Figure 1: Image of CD19^+^ B cell level populations in a B cell depleted mouse v not depleted mouse.** Showing 2 Synapsin-Cre x GCaMP6s male mice (4-6mo. old). The mouse on the left was treated with IgG control antibodies for 3 weeks, while the mouse on the right was treated with anti-CD20 antibodies for 3 weeks.

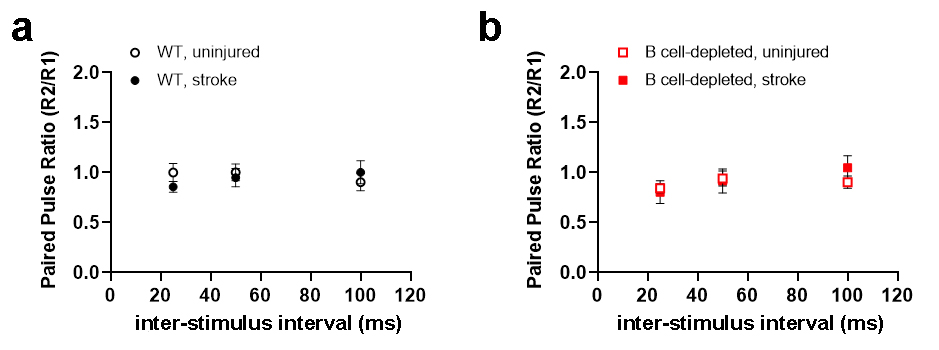

**Supplemental Figure 2:** **Neither stroke nor B cell-depletion alter pre-synaptic release probability of the dentate gyrus**. Paired-pulse ratios of perforant path-DG synaptic transmission (fEPSP slope response 2/response 1) in (*a*) WT (black circles) and (*b*) B cell-depleted mice (red squares) are not different between uninjured (open symbols) and post-stroke (closed symbols) cohort.

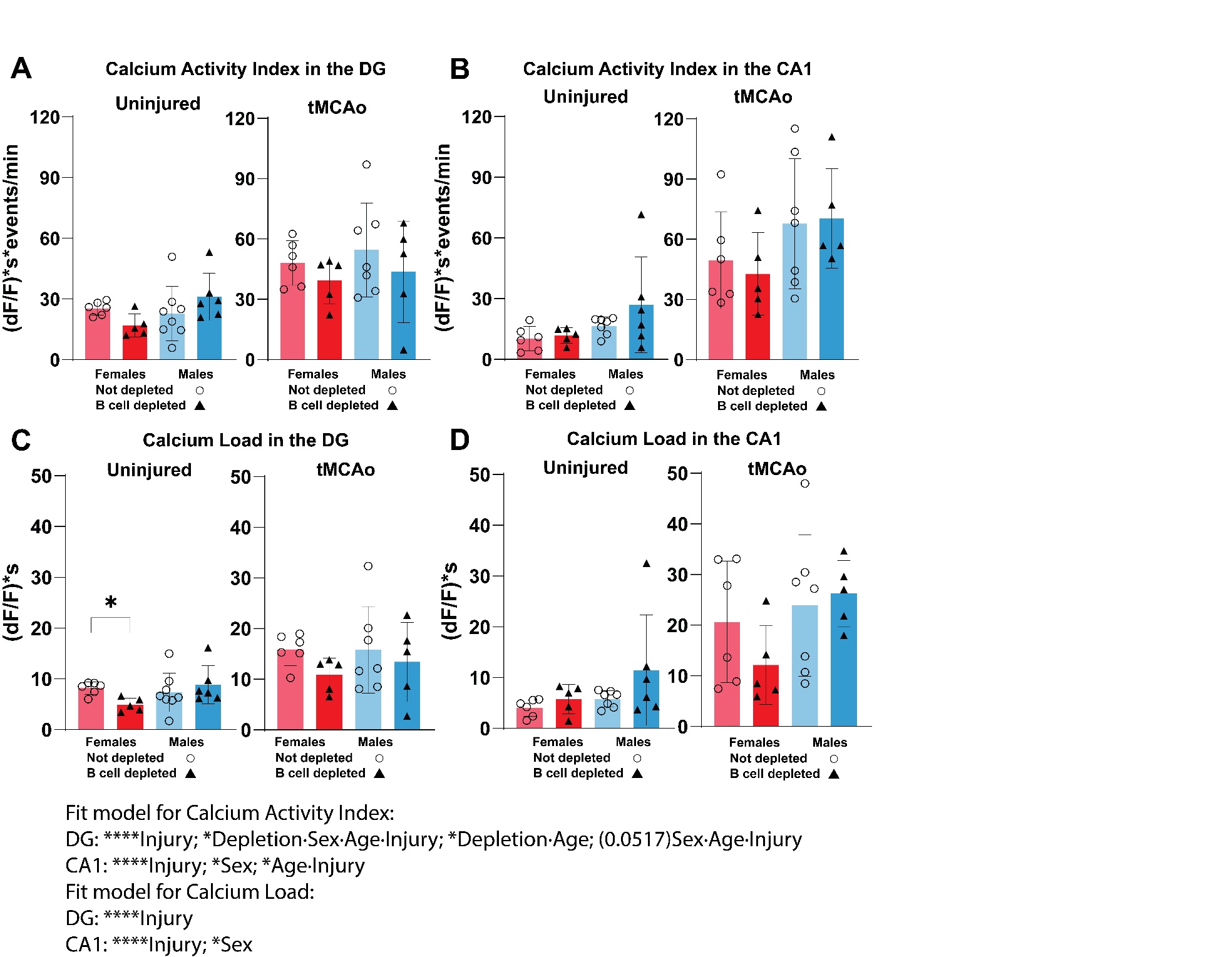

**Supplement Figure 3: Composite Calcium Metrics in CA1 and DG: Activity Index and Calcium Load**

The graphs on the left are for uninjured, right are post-stroke (tMCAo). (A) Calcium Activity Index in the DG. (B) Calcium Activity index in the CA1. (C) Calcium Load in the DG. (D) Calcium Load in the CA1. The bar graph presents means with SD. *p<0.05 as determined using Welch’s ANOVA test, with Dunnett’s T3 multiple comparisons test shown on graph. (A,B,D) Not significant. (C) For Uninjured, Welch’s ANOVA, p=0.0131, with multiple comparison’s shown on graph.

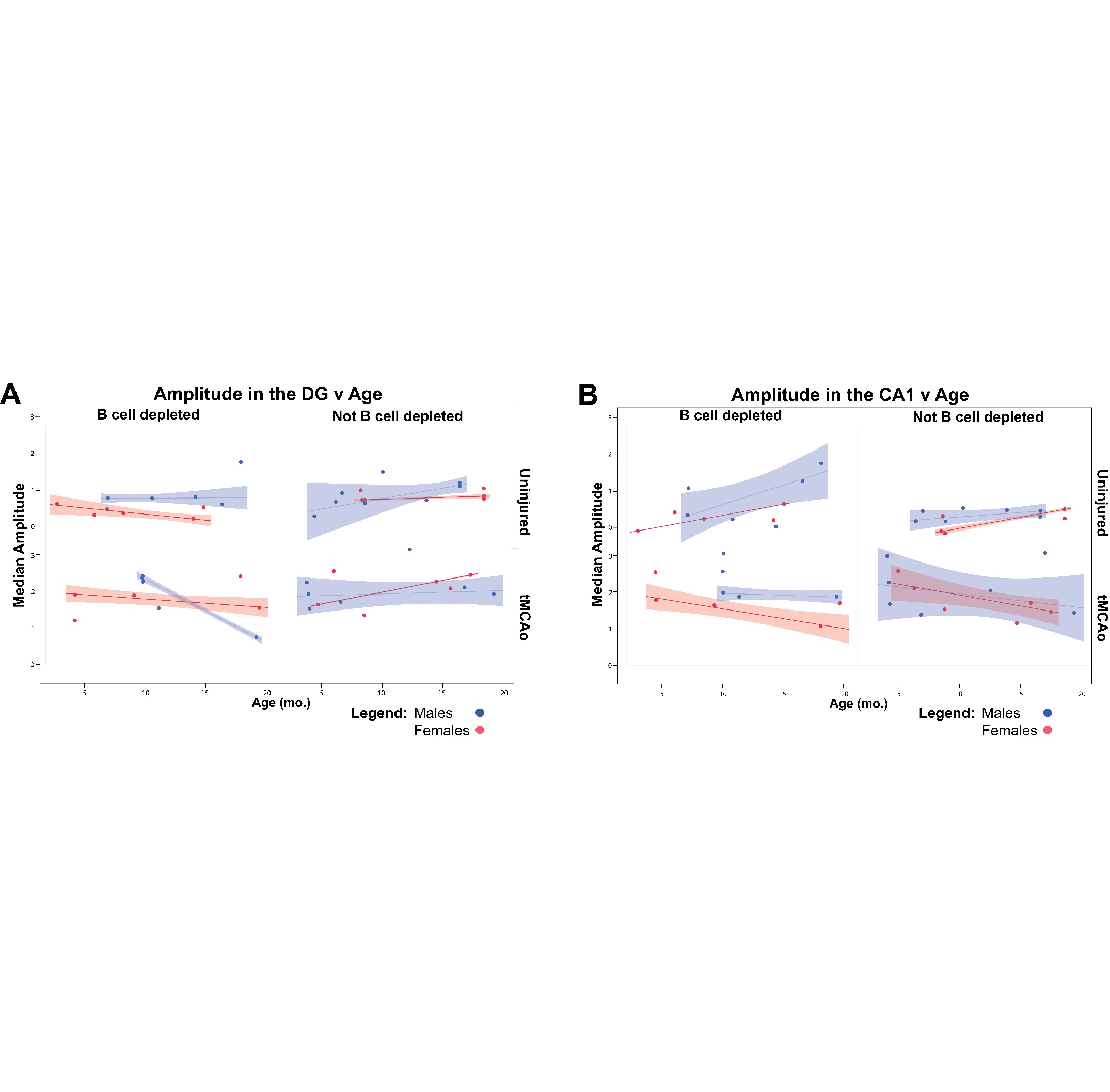

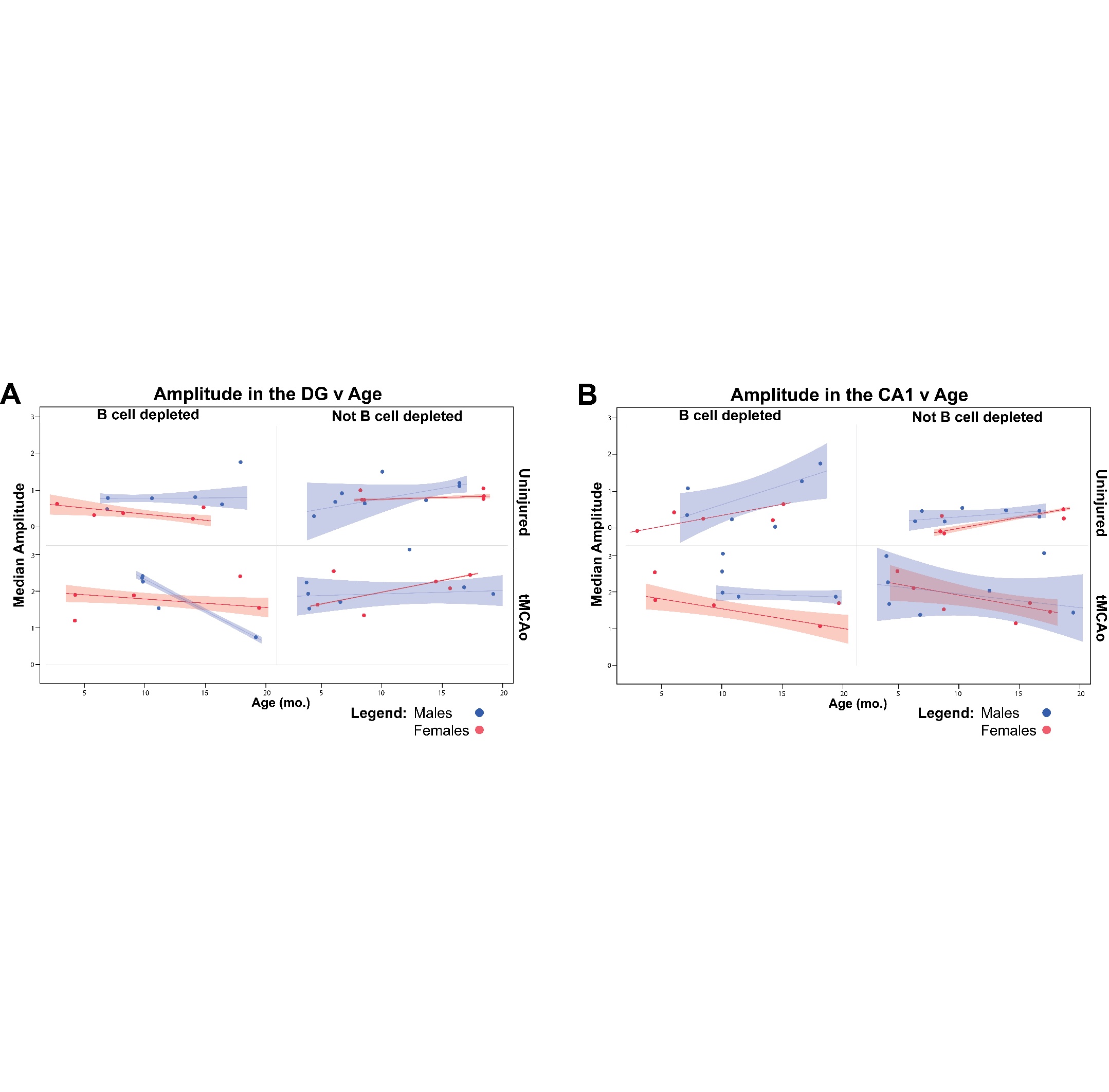

**Supplement Figure 4: Calcium Amplitude versus Age in DG and CA1 across sex, B cell depletion, and uninjured vs tMCAo.**

This figure used a robust regression with the Robust Cauchy method in JMP Pro. This approach applies a weighting scheme that diminished the impact of obersvations with large residuals, prviding more reliable parameter estimates.

A) DG Amplitude v Age

| Condition | Injury Status | Sex | ChiSq | P-value |
| --- | --- | --- | --- | --- |
| B cell depleted | Uninjured | Male | 0.01 | 0.936 |
| B cell depleted | Uninjured | Female | 15.37 | <0.0001 |
| B cell depleted | tMCAo | Male | 1462.31 | <0.001 |
| B cell depleted | tMCAo | Female | 10.88 | 0.0010 |
| Not B cell depleted | Uninjured | Male | 4.02 | 0.045 |
| Not B cell depleted | Uninjured | Female | 14.44 | 0.0001 |
| Not B cell depleted | tMCAo | Male | 0.29 | 0.5929 |
| Not B cell depleted | tMCAo | Female | 4769.13 | <0.0001 |

B) CA1 Amplitude v Age

| Condition | Injury Status | Sex | ChiSq | P-value |
| --- | --- | --- | --- | --- |
| B cell depleted | Uninjured | Male | 10.53 | 0.0012 |
| B cell depleted | Uninjured | Female | 133725.4 | <0.0001 |
| B cell depleted | tMCAo | Male | 1.32 | 0.2501 |
| B cell depleted | tMCAo | Female | 23.75 | <0.0001 |
| Not B cell depleted | Uninjured | Male | 3.60 | 0.0579 |
| Not B cell depleted | Uninjured | Female | 337.13 | <0.0001 |
| Not B cell depleted | tMCAo | Male | 1.11 | 0.2916 |
| Not B cell depleted | tMCAo | Female | 11.52 | 0.0007 |

| Sex | B cell Depletion | Stroke | Mean Age  (mo.) | Mean infarct Size (mm^3^) | Animals | Cell Counts | | |
| --- | --- | --- | --- | --- | --- | --- | --- | --- |
|  |  |  |  |  |  | **CA1** | | **DG** |
| Male | No | No | 10.5 |  | 8 | 885 | | |
|  |  |  |  |  |  | 320 | 565 | |
| Male | Yes | No | 11.9 |  | 6 | 2463 | | |
|  |  |  |  |  |  | 745 | 1718 | |
| Male | No | Yes | 9.6 | 25.18 ± 22.25 | 7 | 1893 | | |
|  |  |  |  |  |  | 906 | | 987 |
| Male | Yes | Yes | 11.7 | 35.18 ± 16.61 | 5 | 1165 | | |
|  |  |  |  |  |  | 691 | | 474 |
| Female | No | No | 13.6 |  | 6 | 512 | | |
|  |  |  |  |  |  | 169 | | 343 |
| Female | Yes | No | 8.8 |  | 5 | 2199 | | |
|  |  |  |  |  |  | 769 | | 1430 |
| Female | No | Yes | 10.7 | 38.99 ± 20.38 | 6 | 1570 | | |
|  |  |  |  |  |  | 822 | | 748 |
| Female | Yes | Yes | 11.3 | 24.04 ± 19.17 | 5 | 1378 | | |
|  |  |  |  |  |  | 768 | | 610 |

**Supplement Table 1: Summary of mice, group characteristics, and cells recorded.** All mice used were Synapsin-Cre x GCAMP6s mice. B cell depletion refers to anti-CD20 treated animals, while ‘No’ referes to IgG control treated animals. ‘Stroke’ refers to animals which received a 45-min tMCAo. Age is shown in months. Infarct size was measured with MRI at 1 week post-tMCAo. Cell counts at the top refer to total cells recorded, and below are split by CA1 region and dentate gyrys (DG) region in the hippocampus. In post-stroke mice, all recording was conducted in the contralesional hippocampus.

| Parameter | Effect | DF | Sum of Squares | F Ratio | Prob > F |
| --- | --- | --- | --- | --- | --- |
| Median (Amplitude, DG) | Injury | 1 | 5.0168 | 48.9517 | <.0001 |
| Median (Amplitude, DG) | Depletion | 1 | 0.8931 | 8.7152 | 0.0059 |
| Median (Amplitude, DG) | Age*Injury | 1 | 0.8001 | 7.8076 | 0.0087 |
| Median (Amplitude, DG) | Depletion*Age | 1 | 1.0089 | 9.8449 | 0.0036 |
| Median (Amplitude, DG) | Sex*Age*Injury | 1 | 2.0078 | 19.5912 | 0.0001 |
| Median (Amplitude, DG) | Depletion*Sex*Age | 1 | 0.4802 | 4.6856 | 0.0380 |
| Median (Amplitude, DG) | Depletion*Age*Injury | 1 | 0.4613 | 4.5018 | 0.0417 |
| Median (Amplitude, DG) | Depletion*Sex*Age*Injury | 1 | 0.7529 | 7.3473 | 0.0107 |
| Median (Events/min, DG) | Depletion*Sex*Age*Injury | 1 | 1.2948 | 5.1069 | 0.0308 |
| Median (Duration, DG) | Depletion*Injury | 1 | 0.3389 | 8.0962 | 0.0077 |

**Supplement Table 2:** **Summary of statistics from the multiple linear regression model in the DG.** Multiple linear regression was performed using the *Fit Model* platform in JMP Pro. A Standard Least Squares linear regression was used to evaluate the influence of each predictor variable on the outcome measures to identify statistically significant effects. Model terms were selected based on experimental design and biological relevance. The outcome variables included amplitude, duration (in milliseconds), and events per minute. Predictor variables included injury (tMCAo vs. uninjured), B cell depletion (depleted vs. IgG control), sex, and age (as a continuous variable in months).

| Parameter | Effect | DF | Sum of Squares | F Ratio | Prob > F |
| --- | --- | --- | --- | --- | --- |
| Median (Amplitude, CA1) | Injury | 1 | 13.4238 | 63.8924 | <.0001 |
| Median (Amplitude, CA1) | Sex | 1 | 1.7234 | 8.2029 | 0.0074 |
| Median (Amplitude, CA1) | Depletion | 1 | 0.8017 | 3.8162 | 0.0598 |
| Median (Amplitude, CA1) | Age*Injury | 1 | 2.1218 | 10.0992 | 0.0034 |
| Median (Events/min, CA1) | Injury | 1 | 3.8053 | 8.6713 | 0.0061 |
| Median (Events/min, CA1) | Depletion*Sex | 1 | 1.7322 | 3.9473 | 0.0558 |
| Median (Duration, CA1) | Sex*Age | 1 | 0.5185 | 5.3679 | 0.0273 |

**Supplement Table 3:** **Summary of statistics from the multiple linear regression model in the CA1.** Multiple linear regression was performed using the *Fit Model* platform in JMP Pro. A Standard Least Squares linear regression was used to evaluate the influence of each predictor variable on the outcome measures to identify statistically significant effects. Model terms were selected based on experimental design and biological relevance. The outcome variables included amplitude, duration (in milliseconds), and events per minute. Predictor variables included injury (tMCAo vs. uninjured), B cell depletion (depleted vs. IgG control), sex, and age (as a continuous variable in months).
